# NeighborFinder: an R package inferring local microbial network around a species of interest

**DOI:** 10.64898/2025.12.05.692507

**Authors:** Mathilde Sola, Adrien Paravel, Sandrine Auger, Jean-Marc Chatel, Florian Plaza-Oñate, Emmanuelle Le Chatelier, Marion Leclerc, Patrick Veiga, Clémence Frioux, Mahendra Mariadassou, Magali Berland

## Abstract

1

**Motivation:** Understanding interactions from microbiome data is a central aspect in microbial ecology, as it provides insights into ecosystem stability, disease mechanisms, and can be used to design synthetic communities. Current network inference tools reconstruct global networks from co-abundance data, which means they capture the overall correlation structure for the entire set of taxa considered. These approaches are computationally intensive and suboptimal when the focus is on the local neighborhood of specific taxa of interest.

**Results:** We introduce NeighborFinder, a local network inference method that enables the targeted discovery of direct neighbors around a species of interest. Using cross-validated multiple linear regression with *𝓁*_1_ penalty and microbiome-specific filters, our approach infers interpretable species-centered interactions, with F1 score ≥ 0.95 on simulated cohorts ranging from 250 to 1000 samples. This method is well-suited for large metagenomic datasets and is particularly valuable for exploratory studies where the targeted hypotheses outweigh the need for global community structure. The approach complements existing methods by being a biologically intuitive and computationally efficient.

**Availability and Implementation:** The R package is freely available on GitHub: https://github.com/metagenopolis/NeighborFinder. The data and source code used to calculate performances and produce the use case example in this paper can be found respectively at: https://doi.org/10.57745/UPITJ0 and https://doi.org/10.57745/HJLWW4.

**Supplementary information:** Supplementary data are available

## 2 Introduction

In recent years, metagenomic sequencing has revolutionized our understanding of microbial communities and enabled researchers to identify species, estimate their abundances, and infer functional potential. As metagenomic datasets grow in scale and resolution, so does the interest in moving towards a more mechanistic understanding of microbial communities. Microbial networks, where the nodes are taxa and the edges represent total or partial correlations - that can be interpreted as potential ecological associations - offer a powerful framework to explore the structure of these communities [1].

Existing network inference methods, mostly based on Graphical Gaussian Models (GGM) [2], rely on global modeling approaches and infer the full network, at great cost in terms of computational burden and statistical power. However, researchers may be primarily interested not in the whole network but only in the neighbors of a specific species of interest (*e*.*g*. a pathogen, a probiotic) [3] as a proxy for direct interactions. Those neighbors can for instance support the design of consortia promoting or inhibiting the growth of a species [4].

Here, we introduce NeighborFinder, a local inference method integrated into an R package that enables fast (running time < 1 minute up to 1 100 samples), efficient and scalable identification of direct neighbors around a species of interest.

## 3 Overview of the method

NeighborFinder is tailored to microbiome data. It was specifically developed for shotgun metagenomic data and includes a default normalization step for such datasets, but can accommodate metabarcoding data (and other count-based inputs) by skipping this step.

The primary function for inferring the local neighborhood of a species is `apply_neighborfinder()`. The R package provides helper functions to visualize results, either as a network or a table, and to construct consensus based on the intersection of multiple inferences. A [technical report](https://metagenopolis.github.io/NeighborFinder/articles/NeighborFinder_technical_report.html) and a [vignette](https://metagenopolis.github.io/NeighborFinder/articles/NeighborFinder_vignette.html) illustrate typical use cases and provide methodological details.

Here, we show how NeighborFinder infers local networks from bacterial abundances in shotgun metagenomic data, focusing on the neighborhoods of three well-known species.

### 3.1 NeighborFinder description

NeighborFinder performs a three-step procedure illustrated in Fig. 1A and detailed below.

1. **Data preparation**. Input data is a species-by-sample abundance table. The first step filters out low-prevalence (*≤* `prev level`) species, for which network interaction detection is notoriously difficult, especially with limited sample sizes [5]. Next, shotgun metagenomic abundances are transformed into count-like values (with `get_count_table()`), by scaling and rounding all values such that the minimum non-zero count is 1. Finally, `mclr` normalization is applied to the previously transformed abundance data. This adapted version of the `clr` normalization preserves the zeros in the dataset, doesn’t require the addition of pseudo counts, and is used in network inference [6].
2. **Network inference**. The local species abundance is regressed against all other using *𝓁*_1_ penalized linear regression and neighbors are species with non-null regression coefficients, following [3]. The regression is performed using `glmnet::cv.glmnet()` [7], with cross-validation used to select the *𝓁*_1_ penalty parameter *λ*. The regression is applied 10 times, with 10 different Random Number Generator (RNG) seeds and the results are filtered in each run to keep only the topmost coefficients in absolute value (`top_filtering` = 30% by default) and limit spurious detection of neighbors.
3. **Network stabilization**. To further improve robustness and reproducibility, only neighbors consistently detected in at least half of the runs are kept. For the conserved edges, the final coefficient is computed as the median over all runs.

**Figure 1.**
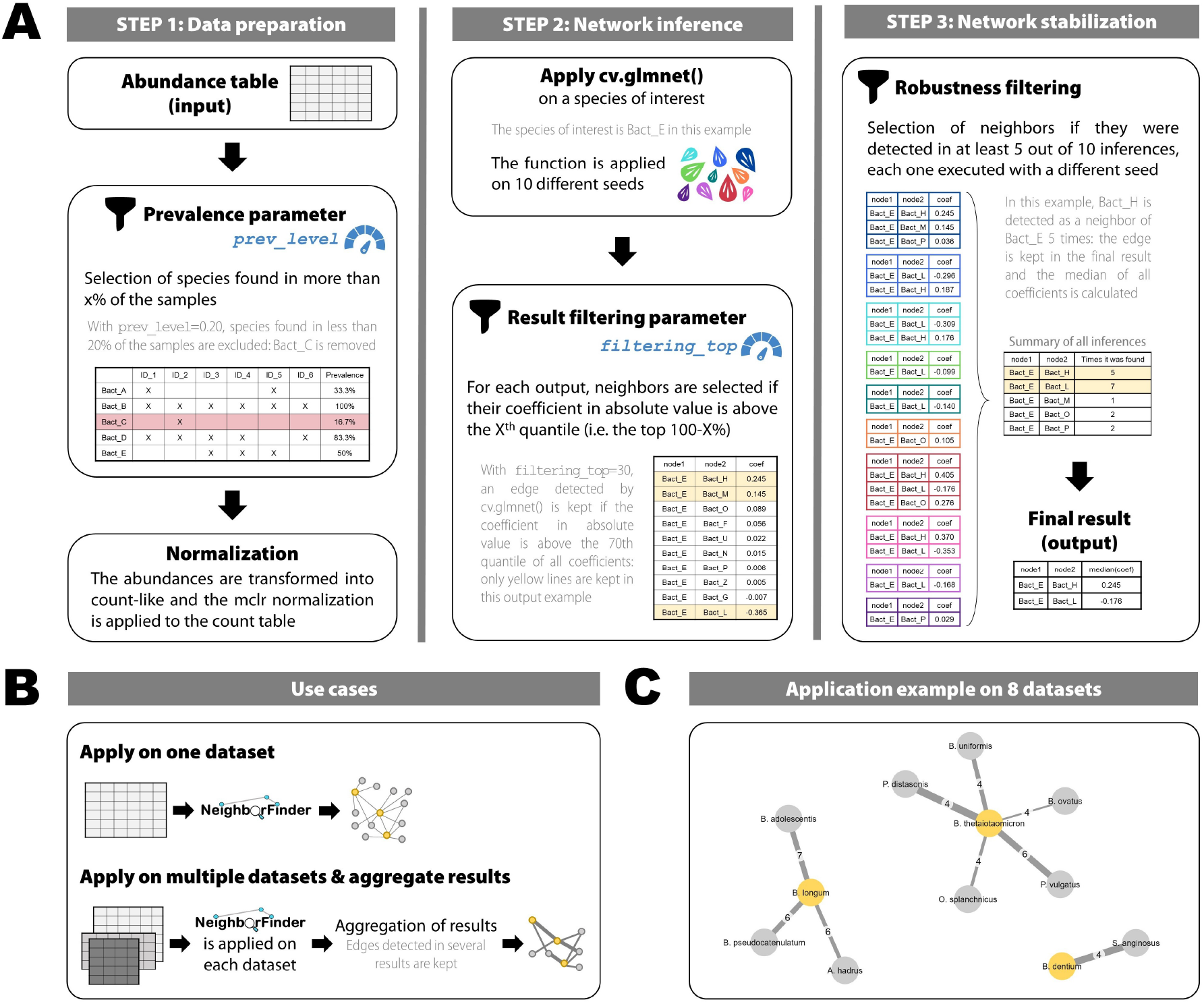
Overview of NeighborFinder, its use cases and an example. A) Presentation of the key function `apply_neighborfinder()` with metagenomic data as input (microbial abundances for each sample), highlighting the two parameters that can be adjusted by the user: `prev_level` and `filtering_top`. B) Description of two main NeighborFinder use cases: on a single dataset, or on multiple datasets for aggregation purposes. C) Aggregated network of an application example focused on three species of interest (yellow nodes): *Bifidobacterium longum, Bifidobacterium dentium*, and *Bacteroides thetaiotaomicron* in eight datasets. The edge label is the number of datasets in which the edge was detected ; the width is proportional to the mean coefficient value.

### 3.2 Performance of the method

#### 3.2.1. Simulation strategy

The function `apply_neighborfinder()` has two key parameters (`prev_level` and `top_filtering` shown in Fig. 1A) that shape the output. It is also well documented that the dataset size has an impact on the inferred network as more samples lead to more edges [2].

Our performance assessment strategy therefore consisted in covering a range of values for both parameters for sizes observed in the literature, with the aim of providing users with guidelines to pick the most suitable combination depending on their dataset size.

We then tested the performance of NeighborFinder on eight independent shotgun metagenomic cohorts (sample sizes from *n*=347 to *n*=1084). For each cohort, similar microbiome data was simulated with known interaction networks (see Supplementary Material). NeighborFinder’s performance was evaluated using precision and recall, aggregated into the F1 score (harmonic mean of recall and precision).

#### 3.2.2 Evaluation procedure

For each bacterial species, local networks were inferred using `prev_level` values between 15% and 35% (in 5% increments). Each setting was repeated 10 times with different RNG seeds, for all simulated datasets. The `glmnet::cv.glmnet()` function was applied, using `cvglm_to_coeffs_by_object()` from the NeighborFinder package. The resulting inferences were then filtered following the procedure implemented in `apply_NeighborFinder()` keeping only the top `filtering_top`% coefficients, ranging from 5% to 30% (in 5% increments), and including 100% to assess unfiltered results (see Fig. 1A, steps 1 and 2).

Finally, for each species, a network was obtained through the stabilization step implemented in NeighborFinder (see Fig. 1A step 3), where edges were retained if detected in at least 5 of the 10 runs across different seeds. F1-scores were calculated for each species, then averaged across species and cohorts.

### 3.3 Determining the best combination of parameters

Figure S1 summarizes performance scores calculated on simulated dataset with sizes ranging from n=100 to n=1000, and Figure S2 focuses on n=50 using a wider range of prevalence filters. They show that parameters `prev_level` and `filtering_top` should be adjusted according to the size of the dataset. When *n* is large, many parameter combinations lead to similar performance (dark green area, F1 score *≥* 0.9 in Figure S1). The best strategy is to select a low value of `prev_filter`, to include as many species as possible, before adjusting `filtering_top` value to ensure good performance (*e*.*g*. with n=1000, prev level=0.15 and filtering top=30). As *n* decreases, increasing `prev_level` while keeping a high`filtering_top` value is required to maintain high F1-scores. In particular, for small datasets (*n*=50), Figure S2 highlights that stringent prevalence filtering (`prev_level` *≥* 0.4) is mandatory to achieve a F1 score above 0.7.

## 4 Usage scenario

### 4.1 Use cases

The method can be applied to one dataset or to several independent datasets (Fig. 1B). In the latter case, NeighborFinder is applied to each dataset and a consensus is built from edges detected across several datasets (*e*.*g*. at least 3 out of 5 datasets) to enhance the overall robustness of the inferred neighborhood. This strategy is illustrated in Fig. 1C, with details in Table 1.

**Table 1.** Application example listing detected neighbors (Node 2) of 3 species of interest (Node 1), with an aggregation strategy. For each edge, the occurrence in the 8 datasets, the mean coefficient (Mean coef) and standard deviation (SD) are indicated along with a potential biological interpretation, supported by references.

| Node 1 | Node 2 | Occurrence | Mean coef | SD | Biological interpretation |
| --- | --- | --- | --- | --- | --- |
| *Bacteroides thetaiotaomicron* | *Bacteroides ovatus* | 4/8 | 0.054 | 0.016 | Functional complementarity [8] |
| *Bacteroides thetaiotaomicron* | *Parabacteroides distasonis* | 4/8 | 0.21 | 0.092 | Taxonomic proximity, niche preference [5] |
| *Bacteroides thetaiotaomicron* | *Bacteroides uniformis* | 4/8 | 0.13 | 0.084 | Taxonomic proximity, niche preference [5] |
| *Bacteroides thetaiotaomicron* | *Odoribacter splanchnicus* | 4/8 | 0.090 | 0.054 | Potential cross-feeding |
| *Bacteroides thetaiotaomicron* | *Phocaeicola vulgatus* | 6/8 | 0.18 | 0.082 | Functional complementarity [9] |
| *Bifidobacterium longum* | *Anaerostipes hadrus* | 6/8 | 0.11 | 0.053 | Potential cross-feeding [10] |
| *Bifidobacterium longum* | *Bifidobacterium pseudocatenulatum* | 6/8 | 0.15 | 0.040 | Taxonomic proximity, niche preference [5] |
| *Bifidobacterium longum* | *Bifidobacterium adolescentis* | 7/8 | 0.15 | 0.058 | Taxonomic proximity, niche preference [5] |
| *Bifidobacterium dentium* | *Streptococcus anginosus* | 4/8 | 0.20 | 0.060 | Niche preference [11] |

### 4.2 Application example

NeighborFinder was applied to eight independent human gut microbiome datasets (details of each dataset in Supplementary Material) to identify the neighbors of three species: *Bifidobacterium longum* (probiotic), *Bifidobacterium dentium* (opportunistic pathogen), and *Bacteroides thetaiotaomicron* (gut symbiont). The eight datasets are used to find robust and ubiquitous neighbors (detected in at least 4 out of the 8 datasets). Fig. 1C shows the resulting network and Table 1 lists the following descriptors for each edge: occurrence (number of datasets where detected), mean coefficient and standard deviation, and biological interpretation of the edge based on literature review.

In particular, the interaction between *B. thetaiotaomicron* and *Bacteroides ovatus* may reflect functional complementarity: *B. ovatus* harbors different glycoside hydrolases and polysaccharide utilization loci (PULs) able to degrade complex plant polysaccharides such as hemicelluloses (galactomannans and xylans) that *B. thetaiotaomicron* lacks [8]. Such complementarity characterizes *B. ovatus* as a companion species to *B. thetaiotaomicron*. Similarly, *B. vulgatus* complements the enzymatic activity of

*B. thetaiotaomicron* [9]. The interaction between *B. longum* and *Anaerostipes hadrus* may plausibly involve cross-feeding, as previously demonstrated between *Bifidobacterium* species and butyrate-producing bacteria such as *Anaerostipes* species [10]. Other detected neighbors may be explained by taxonomic proximity or niche similarity, as often happening with edges detected by network inference methods when the environment is not explicitly accounted for [5]. For example, *B. dentium* and *Streptococcus anginosus* both inhabit the human oral cavity [11].

Although they have different goals, we compared our findings to SPIEC-EASI [12]. It was applied to each dataset and a consensus network was built from the eight inferred networks before extracting the neighbors of the three previous species (see Supplementary Materials). SPIEC-EASI was 37 times slower (4-core parallelization) than NeighborFinder (single core), as expected for the reconstruction of the full network, and missed 6 of the 9 edges found by NeighborFinder, including all neighbors of *B. thetaiotaomicron*, due to lower statistical power for the specific neighborhood reconstruction problem. This application illustrates the variety of biological insights that can emerge from co-abundance patterns in NeighborFinder local networks and its advantage over global network methods.

## 5 Discussion

NeighborFinder provides an efficient and interpretable approach complementary to global network inference. This framework is particularly suited for large-scale datasets and exploratory analyses where speed (see Figure S3 for execution time details), simplicity, and species-centered hypotheses are key.

While it addresses key challenges such as compositionality and sparsity through `mclr` normalization and prevalence filtering to exclude low-prevalence taxa, NeighborFinder suffers from the linearity assumption inherent to GGM-based approaches, which may limit the description of certain types of biological interactions (amensalism, parasitism, etc.) [5]. Similarly to global approaches, our method detects relatively few negative edges. These could be interpreted as signals of competition or inhibition between taxa.

To mitigate instability and noise inherent to all network inference methods, we extended the naive `cv.glmnet` approach with additional features specifically tailored to microbiome data. We used *𝓁*_1_ penalized regression combined with data-specific filtering steps, applied post-processing to the results, and introduced the use of multiple repetitions alongside cross-validation to improve robustness.

Although NeighborFinder has been designed for microbiome data, this method could be useful to explore local networks in other context of tabular data ranging from different taxonomic levels to functional modules for example.

