## Supplementary Material for "NeighborFinder: an R package inferring local microbial network around a species of interest"

Mathilde Sola[1,2], Adrien Paravel[3], Sandrine Auger[3], Jean-Marc Chatel[3], Florian Plaza Oñate[1], Emmanuelle Le Chatelier[1], Marion Leclerc[1,5], Patrick Veiga[1,3], Clémence Frioux[2], Mahendra Mariadassou[4,*], Magali Berland[1,*]

[1]Université Paris-Saclay, INRAE, MGP, 78350, Jouy-en-Josas, France
[2]Inria, University of Bordeaux, INRAE, 33400, Talence, France
[3]Université Paris-Saclay, INRAE, MICALIS, 78350, Jouy-en-Josas, France
[4]Université Paris-Saclay, INRAE, MaIAGE, 78350, Jouy-en-Josas, France
[5]Université Clermont Auvergne, INRAE, UMR454 MEDIS, 63000, Clermont-Ferrand, France
* Corresponding authors

#### 1 Datasets used for the NeighborFinder performance evaluation and the illustration example

##### 1.1 Summary of the selected datasets

The following datasets were retrieved. Associated data and metadata are available at `https://doi.org/10.57745/UPITJO`.

Table S1: **Characteristics of the datasets used in this article.**

| Dataset | Project ID | Number of samples | Countries | Clinical status |
|---|---|---|---|---|
| Asnicar et al. [1] | PRJEB39223 | 1084 | USA,GBR | Healthy |
| Vieira-Silva et al. [2] | PRJEB37249 | 842 | FRA,DEU,DNK | Healthy & Diseased |
| Zeevi et al. [3] | PRJEB11532 | 820 | ISR | Healthy |
| Wallen et al. [4] | PRJNA834801 | 667 | USA | Healthy & Diseased |
| Yachida et al. [5] | PRJDB4176 | 597 | JPN | Healthy & Diseased |
| Schirmer et al. [6] | PRJNA319574 | 382 | NLD | Healthy |
| Wang et al. [7] | PRJNA530339 | 351 | CHN | Healthy |
| Jie et al. [8] | PRJEB21528 | 346 | CHN | Healthy & Diseased |

##### 1.2 Dataset inclusion criteria

The following criteria were considered for the choice of cohorts.

1. Criteria on the study:
   - Human gut / stool samples
   - Shotgun metagenomic sequencing
   - Sample size of n > 300 at one given time-point
   - Collected in a Western or industrialized country

- Data and minimal metadata (*i.e.* age, sex, Body Mass Index, and health condition) available

2. Criteria on individuals:
   - Adults donors ($\geq$ 18 years old)

#### 1.3 Data processing

Data was processed separately for each study according to the following procedure.

##### 1.3.1 Data download

Whole Metagenome Sequencing data was downloaded from the European Nucleotide Archive (ENA).

##### 1.3.2 Quality control

All DNA sequencing reads were quality trimmed and filtered from sequencing adapters using fastp [9] (v0.23.2). Remaining contamination by the host genome was removed by aligning the reads against the human reference genome T2T-CHM13v2.0 with Bowtie2 [10] (v2.5.1) and using at least 90% nucleotide identity threshold for filtering, employing samtools [11] (v1.9).

##### 1.3.3 Microbial species annotation

For all samples, genes and microbial species were identified and quantified with METEOR 2 (`https://github.com/metagenopolis/meteor`) [12] using human gut microbial gene catalogue (IGC2, comprising 10.4 million genes), with GTDB R226 release for taxonomic annotation.

##### 1.3.4 Metadata download & curation

Raw metadata were collected from ENA, NCBI, and supplementary materials of the original publications. Variables were standardized across cohorts, and a common set of minimal metadata was retained (age, gender, body mass index (BMI), health status, country, study name).

#### 1.4 Metadata additional flagging

To obtain the final dataset used in this article, samples were excluded on the criteria detailed below. They follow these reasons: cross-sample contamination, low sequencing depth, longitudinal, missing metadata, and antibiotics use. To do this, samples with a value of "yes" in the "flag" column of the metadata table were removed.

##### 1.4.1 Cross-sample contamination identification

Cross-sample contamination was checked with CroCoDeEL (v1.0.8, `https://github.com/metagenopolis/CroCoDeEL`) [13]. Samples were excluded if species added by contamination exceeded 12 % or if the contamination rate was $>$ 1 % with additional species exceeding 10 %.

##### 1.4.2 Longitudinal series

Samples originating from longitudinal studies (time points beyond baseline, *i.e.* $t > 1$) were flagged.

##### 1.4.3 Missing information on the clinical status

Samples with missing health status were flagged.

##### 1.4.4 Antibiotics consumption

Samples indicating a current antibiotic use were flagged.

#### 1.5 Additional filtering

Samples with a sequencing depth $<$ 20,000,000 reads (paired or single) were excluded.

### 2 Performance evaluation procedure

The performance of a network inference method can only be rigorously assessed through data simulation. By generating synthetic datasets derived from real ones, we obtain a defined ground truth that allows us to evaluate whether the method recovers the expected results, while maintaining key structural features of the data, such as sparsity. Simulation further allows us to vary the number of samples (e.g., from 50 to 1000), even when the original dataset contains only 300 samples for example.

Hence, we simulated data with known structure based on each cohort characteristics (see below). We applied NeighborFinder on each simulated dataset with different values of `prev_level` and `filtering_top` parameters. We then checked the inferred results to compare them with the original structure and calculated performance metrics. This was done for several samples sizes to explore the span of the method's performance.

NeighborFinder's performance was evaluated using the F1 score (harmonic mean of recall and precision) and averaged on the 8 simulated datasets (Figures S1 and S2).

#### 2.1 Data simulation

For each of the eight datasets mentioned above, a graph with a "cluster-like" structure was generated with the `graph_step()` function. A precision matrix $\Omega$ with non-null coefficients respecting the graph topology and sparsity was produced and then inverted to produce a covariance matrix $\Sigma$.

Semi-synthetic simulated datasets of sizes n=50, n=100, n=250, n=500, and n=1000 samples were generated with `new_synth_data()` using gaussian copula from the covariance matrix $\Sigma$ and the original count matrix to produce count tables that (i) have the same marginal counts distributions as the original cohort (ii) while enforcing the correlation between taxa encoded in $\Sigma$. The graph edges are here considered as true edges (ground truth).

#### 2.2 Performance metrics

Precision, recall and F1 scores were used as performance metrics.

- Precision: $P = \frac{TP}{TP+FP}$
- Recall: $R = \frac{TP}{TP+FN}$ with TP: true positives, FP: false positives, and FN: false negatives
- $F1\_score = \frac{2.P.R}{P+R}$ with P: precision and R: recall

### 3 Time of execution estimation procedure

The execution time of the function `apply_NeighborFinder()` was observed to be impacted by the number of observations (sample size) and the number of variables (impacted by the `prev_level` parameter). We selected the 7 species with at least 90% of prevalence in all 8 datasets. We then applied the function to each dataset (to assess for the size effect) with several values of `prev_level` (15%, 50%, and 80%). All elapsed running times are plotted in Figure S3.

### 4 Comparison with SPIEC-EASI

To see if a global network inference method was able to reconstruct the same neighborhoods, we conducted a comparison under identical conditions with SPIEC-EASI [14]. We applied SPIEC-EASI (with `rep.num = 100` and 4-core parallelization) to each of the eight datasets used in Fig. 1C, reconstructing full global networks. This process required 5 hours, 39 minutes, and 23 seconds total (details on machine configuration below). In contrast, NeighborFinder reconstructed all eight local neighborhoods (centered on *B. thetaiotaomicron*, *B. longum*, and *B. dentium*) in just 8 minutes and 51 seconds, without any parallelization.

To enable fair comparison, we extracted only the edges incident to the three target species from each SPIEC-EASI full network and applied identical aggregation and thresholding criteria (occurrence $\geq$ 4) as for NeighborFinder.

As shown in Figure S4, SPIEC-EASI recovered 3 out of the 9 high-confidence edges consistently detected by NeighborFinder—namely, those also observed in our original analysis—but with lower or equal

detection frequency across datasets. Notably, SPIEC-EASI identified one additional edge between *B. dentium* and *Streptococcus vestibularis*, which was detected in only a single dataset by NeighborFinder. However, SPIEC-EASI failed to recover six interactions detected by NeighborFinder (Table 1), all of which are supported by the literature with four independent sources, including the entire neighborhood surrounding *B. thetaiotaomicron* and an additional interaction involving *B. longum*.
The code used to compute this comparison is on the repository, together with the cohorts `https://doi.org/10.57745/HJLWW4`. Note that SPIEC-EASI was run using parallelization on 4 cores of a AMD EPYC 9554 - 3.10 GHz / 3.75 GHz (Turbo) server, while the NeighborFinder job ran on much slower Intel Xeon E5-4600 (2.4 Ghz) core.

### 5 Supplementary Figures

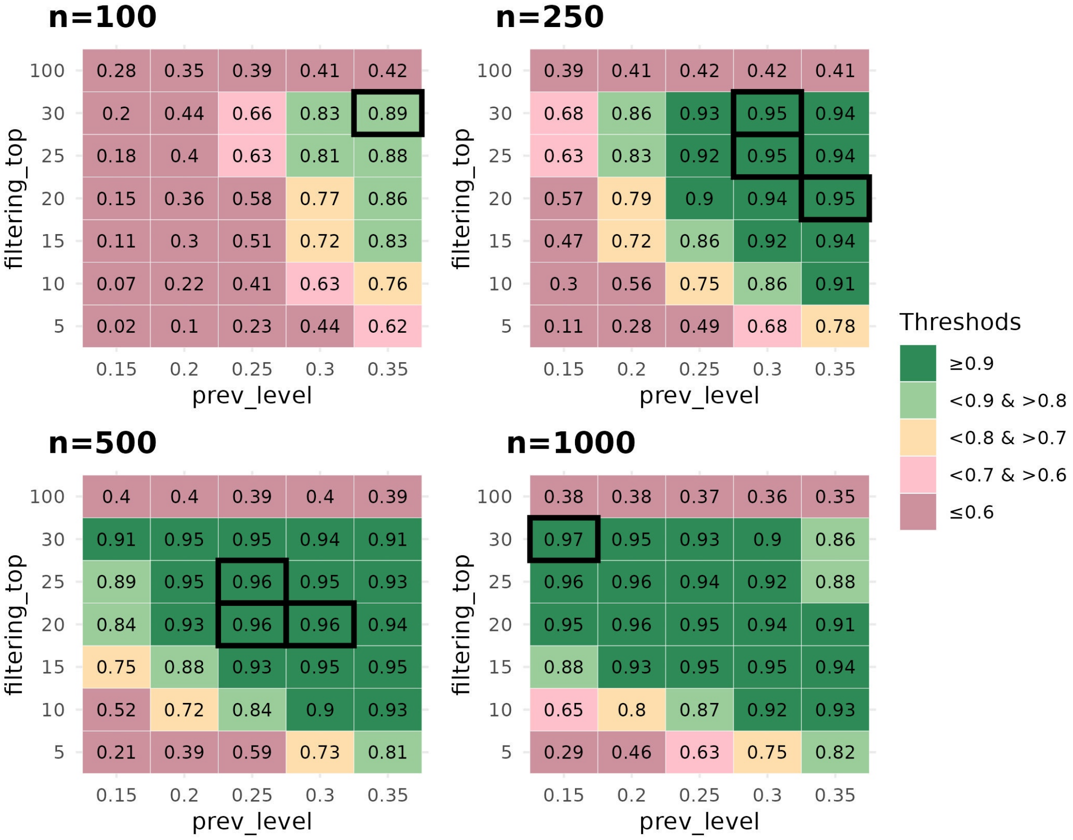


Figure S1: **NeighborFinder performance scores averaged on 8 semi-synthetic simulated data for different values of the function parameters.** The color thresholds indicate a range of parameter combinations that lead to equivalent method performance. The first line (`filtering_top`=100) corresponds to the naive behaviour of `cv.glmnet()`, where end of step 2 and step 3 are not applied (see Fig. 1A).

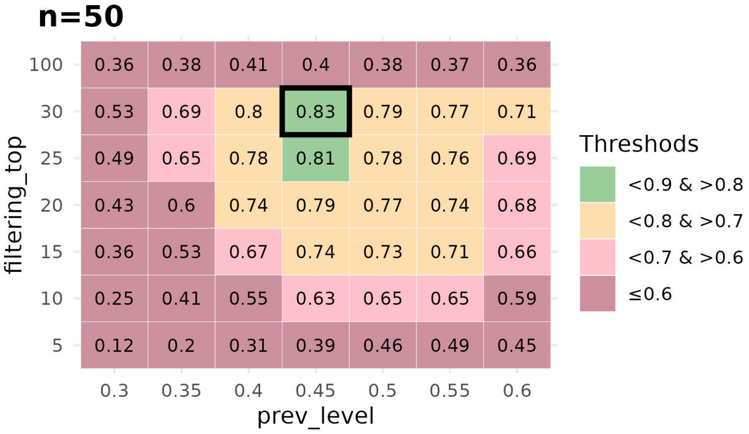


Figure S2: **NeighborFinder performance scores averaged on 8 semi-synthetic simulated data of size n=50 for different values of the function parameters.** The `prev_level` parameter range includes more values. The color thresholds indicate a range of parameter combinations that lead to equivalent method performance. The first line (`filtering_top`=100) corresponds to the naive behaviour of `cv.glmnet()`, where end of step 2 and step 3 are not applied (see Fig. 1A).

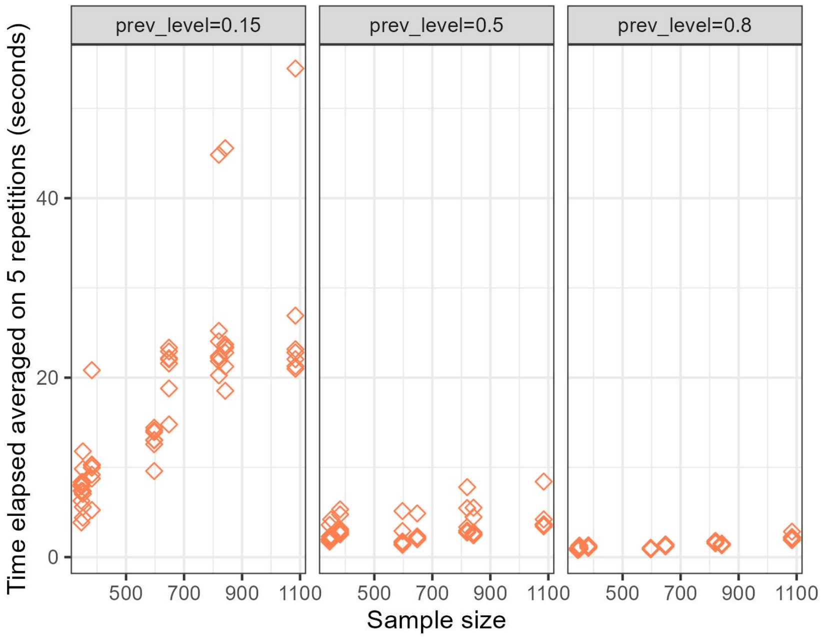


Figure S3: **NeighborFinder execution time for several `prev_level` values and dataset sizes.** Each point is an execution time averaged on 5 repetitions.

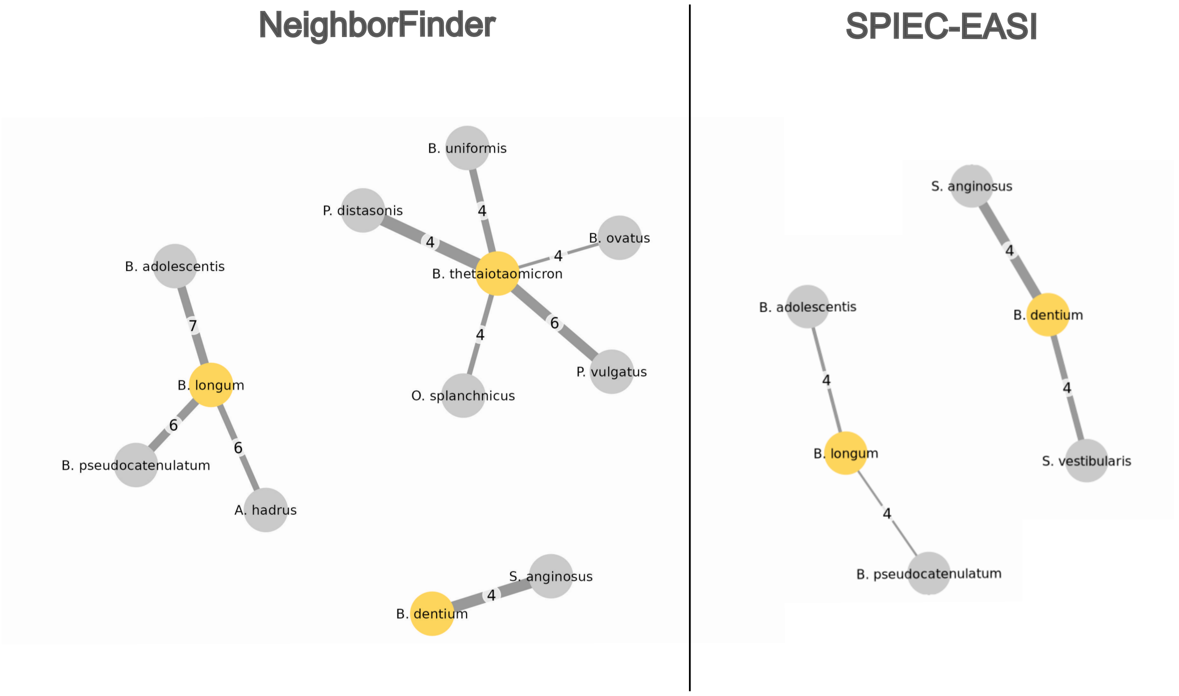


Figure S4: **Comparison of aggregated networks generated by NeighborFinder and SPIEC-EASI.** The comparison was done on the application example, focusing on three species of interest (yellow nodes), with eight independent datasets. The edge label is the number of datasets in which the edge was detected ; the width is proportional to the mean coefficient value.

[13] Lindsay Goulet, Florian Plaza Oñate, Alexandre Famechon, Benoît Quinquis, Eugeni Belda, Edi Prifti, Emmanuelle Le Chatelier, and Guillaume Gautreau. CroCoDeEL: accurate control-free detection of cross-sample contamination in metagenomic data, February 2025. Pages: 2025.01.15.633153 Section: New Results.

[14] Zachary D. Kurtz, Christian L. Müller, Emily R. Miraldi, Dan R. Littman, Martin J. Blaser, and Richard A. Bonneau. Sparse and compositionally robust inference of microbial ecological networks. *PLOS Computational Biology*, 11(5):1–25, 05 2015.
